# Adolescent Seizure Impacts Oligodendrocyte Development, Neuronal-Glial Circuit Formation, and Myelination

**DOI:** 10.1101/2024.04.05.588329

**Authors:** Kylie Foutch, Iris Tilton, Aundrea Cooney, Robert Brock, Raghav Gupta, Cole Bender, Chloe Franzia, Mary Francis Garcia, Jeni Deneen, Megan Baldemor, Christopher Arellano Reyes, Daniela Moura, Hirofumi Noguchi, Laura Cocas

## Abstract

Myelin sheaths, formed by oligodendrocyte cells in the CNS, are vital for rapid conduction of electrical signals down neuronal axons. Oligodendrocyte progenitors differentiate and myelinate axons during development and following demyelinating injury. However, the mechanisms that drive the timing and specificity of developmental myelination are not well understood. It is known that oligodendrocyte progenitors receive synapses from neurons, providing a potential mechanism for neuronal-glial communication. We have previously shown that changing neuronal activity affects the proliferation of oligodendrocyte cells and neuron to OPC connections. We hypothesized that OPC proliferation and differentiation would be affected by pathological neuronal activity during adolescent development, when developmental myelination is occurring, and that this would also impact neuron to OPC connectivity and myelination. We used kainic acid to induce a seizure, then analyzed changes in the rate of OPC proliferation and differentiation five days later in the cerebral cortex, corpus callosum, and hippocampus. We found that OPC proliferation increased, the overall numbers of OPCs increased, and the number of mature oligodendrocytes decreased. We measured changes in the myelination to determine whether seizure activity directly affected myelination rate in adolescent development, and found decreased myelin in the cerebral cortex, corpus callosum, and hippocampus. We used viral monosynaptic circuit tracing to determine whether connections between neurons and OPCs were affected by seizure activity, and found a decrease in neuron to OPC connections in seizure mice compared to controls. Finally, we measured changes in the presence of kir4.1 potassium channels in OPCs, an important regulator of OPC membrane potential as well as an ion channel important for myelination, and found that there was a decrease in the number of potassium channels on OPCs after adolescent seizure. These findings provide insight into the response of the adolescent brain to seizure activity, as well as how seizures affect neuronal glial connections, OPC development and myelin formation, with the goal of understanding how these mechanisms may be important for treatment of demyelination after seizure and in epilepsy.

## Introduction

Many neurological diseases such as multiple sclerosis (MS) and epilepsy are characterized by maladaptive myelination. Epilepsy is a demyelinating disease that produces periodic, unprovoked seizures and results in demyelination. 50 million people worldwide suffer from epilepsy, with 3.4 million people in the United States alone, and 470,000 children in the U.S. have been diagnosed with active epilepsy (Zack and Kobau, 2015; Russ et al., 2012). Overall, 0.7% of children and adolescents have had a seizure in the previous year (Cui, et al., 2015). Existing treatments manage seizure incidence, but there is still need for the development of effective treatment options that can reverse myelination damage and promote remyelination. Understanding the cellular effects of seizure in the adolescent brain can inform both diagnosis and treatment.

Oligodendrocyte precursor cells (OPCs) make up about 75% of the glial cells in the brain (Pelvig, 2008) and give rise to oligodendrocyte cells (OLs) responsible for myelination. Despite most myelin being produced in the early stages of life, OPCs persist in the adult human brain (Nishiyama et al., 2009). OPCs present diverse morphological and physiological properties and only a small percentage of these cells mature into OLs (Chittajallu et al., 2004). It is well established that neuronal activity contributes to OPC differentiation and myelination (Cavaliere et al., 2012; Fannon et al., 2015; Li et al., 2013; Mitew, et al., 2018; Spitzer et al., 2016; Yuan et al., 1998). Synaptic connections between neurons and OPCs have been observed in the hippocampus (Bergles et al., 2000; Mangin et al., 2008), cerebellum (Lin et al., 2005), and the cochlear nucleus (Müller et al., 2009). However, how seizure-driven pathological neuronal activity affects adolescent OPC development and myelination has not been previously studied. As the only glial cell to receive synaptic input from neurons, OPCs are affected by synaptic activity through alterations to OPC differentiation and proliferation (Bergles et al., 2010). Increasing activity using optogenetic stimulation in pyramidal neurons leads to local increases in OPC proliferation, as well as increased myelin sheath thickness (Chang, Redmond, and Chan, 2016).

Previous work has shown that Wallerian degeneration of axons occurs after kainic acid induced seizure, with loss of myelin within 2-4 days, as well as increased microglia and macrophages (Dusart, Marty, and Peschansky, 1992). Previous work using kainic acid injections focally has found myelin damage 2 days after injection, with vacuoles forming in the CA1 region of the hippocampus (Hopkins et. al., X). A pilocarpine model of status epilepticus in adult rats resulted in a reduction in the quantity of MBP and CC1 in the hippocampus, as well as an increase in OPC markers and OPC proliferation (Lue et al., X). During remyelination after seizure, OPCs increase in number and are the main cells responsible for generating new oligodendrocytes to remyelinate axons (Franklin and Goldman, 2015). Finally, recent work has implicated the Kir4.1 potassium channel, an inwardly rectifying potassium channel, in OPC development, as well as in demyelinating injury after ischemia (Song et al., 2018). However, no one has examined status epilepticus effects on OPC function in adolescent mice, a time period where it is known that neuronal activity is necessary both for synapse development and refinement as well as myelination. In this paper we sought to determine the effects of pathological neural activity via induction of seizure on OPC development, myelination, and connectivity in the adolescent forebrain, examining several key structures affected by seizure: the cerebral cortex, hippocampus, and corpus callosum.

Our results indicated an increase in OPC activity in the hippocampus and cortex and increased OPC proliferation in the somatosensory cortex after kainic acid-induced seizure. We found that in the cerebral cortex, there was an increase in the number of OPCs specifically, while in the corpus callosum, there was a decrease in the total number of oligodendrocyte lineage cells. In the CA1 of the hippocampus, we found a decrease in the number of mature oligodendrocytes, and an increase in the number of OPCs. In the cerebral cortex, corpus callosum, and CA1, we found a decrease in myelination. This was consistent with our circuit tracing data, in which we found a loss in the number of neurons connected to OPCs after seizure. Finally, we found that seizure activity led to a decrease in the number of OPCs expressing the Kir4.1 inward rectifying potassium channel, responsible for 80% of potassium currents in OPCs. These results indicate that the short term effects of seizure affect many aspects of OPC development, circuit formation, and myelination in the developing adolescent brain.

## Methods

### Animals

All animal procedures were approved by the Institutional Animal Care and Use Committees at Santa Clara University and the University of California, San Francisco. Pdgfrα-CreERT2 driver mice were crossed to ROSA-tTA and pTRE-Bi-G-TVA mice and the resulting litters were genotyped in order to generate Pdgfrα-CreERT2; ROSA-tTA; pTRE-Bi-G-TVA mice. (Kang et al., 2010; Han et al., 2013; Yasuda, Mayford., 2006). Following tamoxifen-induced gene recombination, Pdgfrα-CreERT2; ROSA-tTA; TRE-Bi-G-TVA mice express the avian envelope protein TVA and the rabies glycoprotein GRab under tetracycline control, only in OPC cells expressing Pdgfrα. This allows for OPC-specific targeting of the viral proteins necessary for infection (TVA) and spread (GRab). At p28 tamoxifen was delivered at a dose of 0.5mg/40g via IP injection. Saline or Kainic acid was also delivered at his time, as described below in the seizure procedure. At p30 and p33 tamoxifen was again delivered at the above dose. A pseudotyped rabies virus was injected at p35, as described below in the viral circuit tracing methods. Mice were sacrificed via perfusion at p40. These mice were used for the rabies single cell circuit tracing experiments. All other experiments were done on sex-matched littermates. For these littermates, Saline or Kainic acid was injected at p28, as described below in the seizure procedure. Starting at p33, Edu was injected IP for 5 days at a dose of 10mg/kg. Mice were sacrificed via perfusion on p38, 2 hours after their final Edu injection.

### Seizure Induction

Seizures were induced in equal numbers of male and female control mice using consecutive low dose IP injections of kainic acid (KA, Enzo Life Sciences, dissolved in saline (.9%NaCl)) (as in Noguchi et al 2023). The initial KA dose was 5 mg/kg. Every 20 minutes thereafter, the mice received IP injected doses of 2.5 mg/kg KA until an initial Racine stage 4 seizure (rearing with forelimb clonus) or Racine stage 5 seizure (rearing and falling with forelimb clonus, general motor convulsions) was observed. If after receiving a total of 30 mg/kg KA the mice still did not exhibit a stage 4 or stage 5 seizure, these mice were excluded from the experiment. Mice receiving an IP injection of saline were used as the control condition for all experiments.

### Virus

An avian envelope protein pseudotyped (EnvA) G deleted mutant rabies virus that expressed a red fluorescent reporter ((EnvA)SADΔmCherry) was amplified and pseudotyped from stock viruses from the Callaway lab (Salk Institute) following their established protocol (Wickersham, et al, 2010). The virus was titered on mammalian 3T3 cells and 3T3-TVA cells to confirm that the virus was pseudotyped and to determine the viral titer: 1 × 10^9^ IU ml-1 was reached for each round of viral production.

### Viral Circuit Tracing by Brain Region

Pdgfrα-CreERT2; ROSA-tTA; pTRE-Bi-G-TVA (N=4-6/group, counterbalanced for sex) were dosed with tamoxifen (DOSE: 2 mg/25 g body weight) via intraperitoneal (IP) injection at P25, P27, and P29. At P30, mice were stereotaxically injected with 300nl of EnvA pseudotyped G deleted rabies virus expressing mCherry (EnvA ΔGRabV-mCherry, 300nl). We used an attenuated rabies virus that expresses red fluorescent protein, making it possible to analyze retrograde (presynaptic) monosynaptic connections in vivo. By combining attenuated rabies virus with a Cre-loxP based system, we examined neuronal input onto OPCs using the Pdgfrα-CreERT2 driver mouse crossed to ROSA-tTA mice bred to TET-BI-G-TVA mice, inducing recombination in Pdgfrα-expressing OPCs using tamoxifen. These mice then expressed the avian envelope protein TVA and the rabies glycoprotein GRab only in OPCs. This allows us to target only OPCs with the viral proteins necessary for infection (TVA) and spread (GRab). We targeted OPCs with a viral injection of avian envelope protein pseudotyped deletion mutant rabies virus (EnvA) RabV that expresses a red fluorescent reporter (mCherry), prepared in our lab using standard protocols and as we have previously published (Cocas, et al., 2016; Wickersham, et al., 2010). We then stained for immunomarkers of OPCs and examined the neurons (labeled in red) that are presynaptically connected to each OPC population (labeled with both red and green). Whole brains were scanned for imaging and quantified to yield biological replicates for all connected neurons and starter OPCs. We were thus able to quantify the numbers and locations of OPCs and presynaptically connected neurons based on this approach. Using stereotaxic injection to target the CA1 of the hippocampus (X 1.1, Y −1.7, Z 1.4) using a Kopf stereotaxic rig, we infected OPCs for viral circuit tracing. Following injection, mice recovered and were placed in group housing for a period of 5 days to allow for viral expression, after which they were killed and the brains were collected and processed for histology.

### Tissue Processing and Sectioning

Mice were sacrificed for transcardial perfusion under terminal anesthesia with sodium pentobarbital (250 mg/kg). Anesthetized mice were pinned and visualization of the heart was achieved following removal of the rib cage, after which a 21-gauge needle was inserted into the left ventricle, the vena cava snipped, and 10 ml of ice-cold PBS run through the line, followed by 10 ml of 4% paraformaldehyde. After perfusion, the brain was harvested and placed at 4ºC in 4% PFA for 2 hours, after which it was cryoprotected in 30% sucrose overnight. The brain was then embedded in O.C.T. compound (Sakura Finetek; Torrance, CA, USA) on dry ice and stored at −20 ºC until sectioned into 30 μm coronal slices on a CM1860 Leica cryostat (Leica Microsystems). Slices were mounted onto Colorfrost Plus slides (Fisher Scientific) and stored in PBST at 4ºC until ready for staining.

### Immunohistochemistry

Sections were washed thrice with 0.2% PBST for 10 minutes at a time, then incubated in blocking buffer (10% lamb or donkey serum, 0.2% Triton, 0.005% sodium azide, and 1:50000 DAPI in PBS) for 30 minutes, after which they were incubated in primary antibody overnight at room temperature. Following three washes with PBS, sections were then incubated in secondary antibody for 1.5 hours at room temperature, followed by three washes in PBS, and mounted using Fluoromount-G mounting medium (Thermo Fisher Scientific; Waltham, MA, USA). Stained sections were imaged using a Zeiss Axioscanner, and representative images for each condition were collected using a Zeiss 710 confocal microscope. The following primary antibodies were used: anti-Olig2 Rb (Abcam ab109186); anti-Olig2 Ms (Millipore MABN50), anti-CFos Rb (Synaptic Systems 226 008), anti-MBP rat (Millipore MAB386), anti-CC1 mouse (Calbiochem OP80), anti-BCAS rabbit (Bios Antibodies bs-11462R). All secondary antibodies were Alexa Fluor secondary antibodies from Jackson Immunoresearch, raised in goat, and conjugated to 488, 546, or 633 fluorophores as needed.

### EdU Staining

Sections were stained with EdU using the Click-iT® EdU Imaging Kit with Alexa Fluor® 647 Azide (ThermoFisher, C10337, Waltham, MA, USA) according to manufacturer instructions. Following EdU staining, slides were incubated in blocking buffer (10% donkey serum, 0.33% Triton, 0.005% sodium azide, and 1:50000 DAPI in PBS) for 30 minutes, then immunostained overnight with anti-rabbit Olig2. Slides were washed in PBS three times and mounted using Fluoromount-G mounting medium. Representative images for each condition were collected using a Zeiss 710 confocal microscope. The number of EdU+/Olig2+ cells in the hippocampus were quantified for each condition.

### Data Collection and Statistical Analyses

Sections were analyzed in order to manually quantify the number of co-labeled cells by category. For all analyses except the viral circuit tracing experiments, 3 technical replicates were quantified per biological replicate; no images with <2 technical replicates were used. Biological replicates are reported in all figure legends. Zeiss Zen lite was used to visualize the image files from the Zeiss Axioscanner for quantification, and FIJI was used to visualize and count the cells in the image files from the Zeiss 780 Confocal Microscope. All schematics were created with BioRender. For every animal in the viral circuit tracing experiments, we sectioned the entire forebrain in 30μm sections and counted the total number of RabV+ cells without estimation.

Statistical analyses were performed, first testing for normality using the Shapiro-Wilk test for normality. In instances where the distribution of the data was non-Gaussian, non-parametric analyses were completed using the Mann-Whitney test. In instances where the distribution was normal, unpaired t tests were used. Descriptive statistics are included in all figure legends. All data were analyzed and plotted using GraphPad Prism version 8.0.0.

For MBP % area analysis, we used FIJI to threshold all images using the mean threshold of grayscale images before the % area was measured.

## Results

We used a kainic acid model to induce seizure in adolescent mice, with the goal of understanding the impact of a single seizure on the development of oligodendrocytes, neuronal-glial synapses, and developmental myelination at this important developmental stage for synapse formation and myelination. We first confirmed that the kainic acid-induced seizure resulted in increased local activity in the brain regions that have previously been implicated in seizure and epilepsy (Lapato, et al., 2017). Because we were interested in the short term effects of seizure, as opposed to longer term effects after seizure or induction of epilepsy, we sacrificed animals five days after seizure induction at P25, and stained for cfos and Olig2 in the forebrain, analyzing cells in the somatosensory cortex, corpus callosum, CA1 and dentate gyrus of the hippocampus. We found that in all regions there was an increase in the number of c-fos+ cells, as well as c-fos+ Olig2+ cells (Figure 1). This indicated that the kainic acid treatment was sufficient to have a short-term effect on activity, as measured by c-fos labeling, even five days after seizure, and that oligodendrocyte lineage cells also showed increased activation as a result of the seizure.

**Figure 1.**
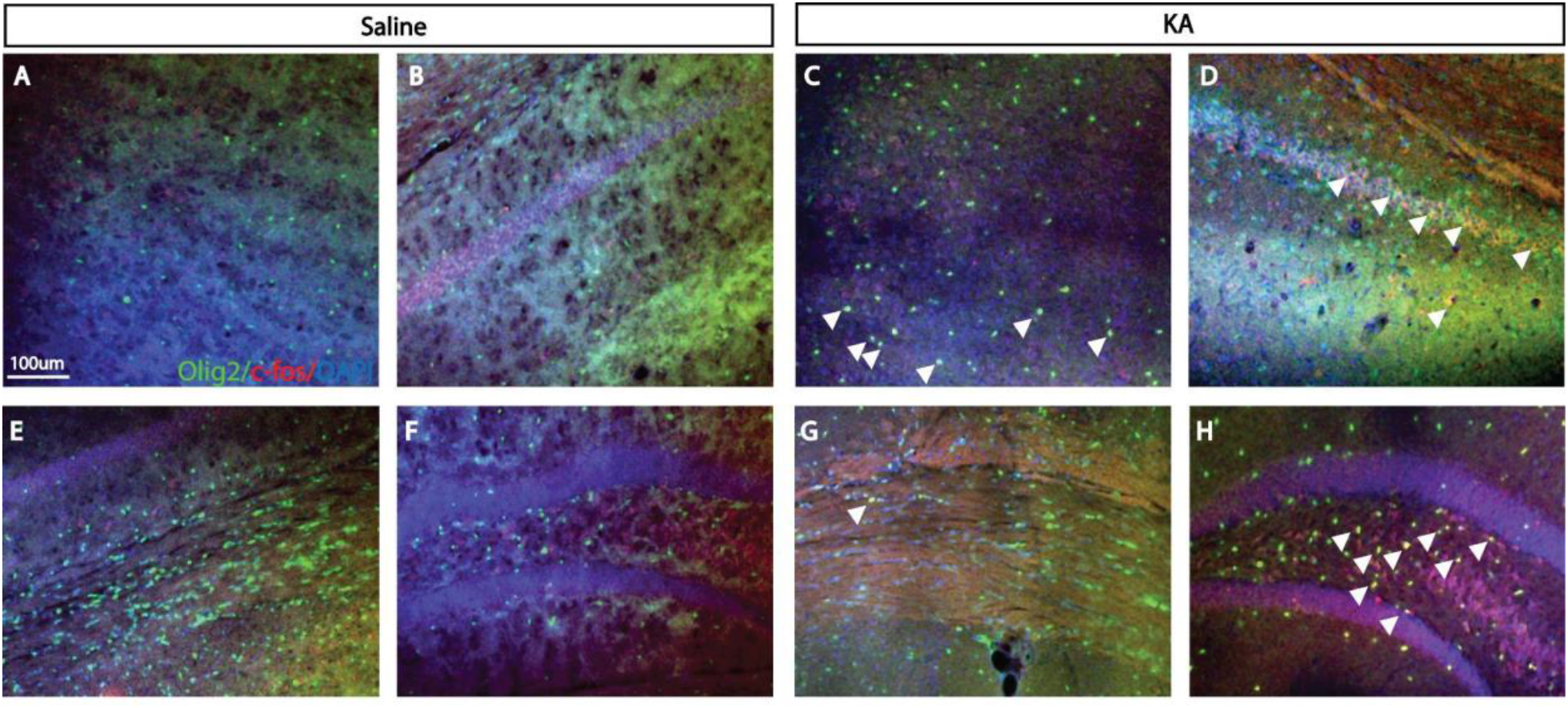
Kainic Acid Induced Seizure Results In Increased Activity in OPCs in the Cortex and Hippocampus in Adolescent Brains. Coronal sections from sex-matched littermates were stained using immunohistochemistry for Olig2 (green) and c-fos (red), and counterstained for DAPI (blue). Control littermates received saline IP injections (A, B, E, and F), and experimental littermates received kainic acid IP injections up to 30 mg/kg (C, D, G, and H). Stains were visualized in the cortex (A and C), CA1 (B and D), corpus callosum (E and G), and DG (F and H). Arrows indicate C-fos+/Olig2+ cells. N=3 (WT); N=3 KA. Scale bar, 100μm.

We next determined the effect of seizure activity on OPC proliferation in the adolescent brain. We injected animals daily with Edu for 5 days, beginning on the day of the seizure. We then used Click-it chemistry to mark Edu-labeled cells, and stained for Olig2, marking all dividing cells from the oligodendrocyte lineage (Figure 2). We found that in the somatosensory cortex there was a significant increase in the percent of Edu+ Olig2+ cells (out of total Olig2+ cells), compared to animals that only received saline, indicating that seizure increased the number of proliferating OPCs in the cerebral cortex. In the corpus callosum, as well as the CA1 and dentate gyrus of the hippocampus, there were not significantly more proliferating OPCs in animals after seizure compared to saline controls.

**Figure 2.**
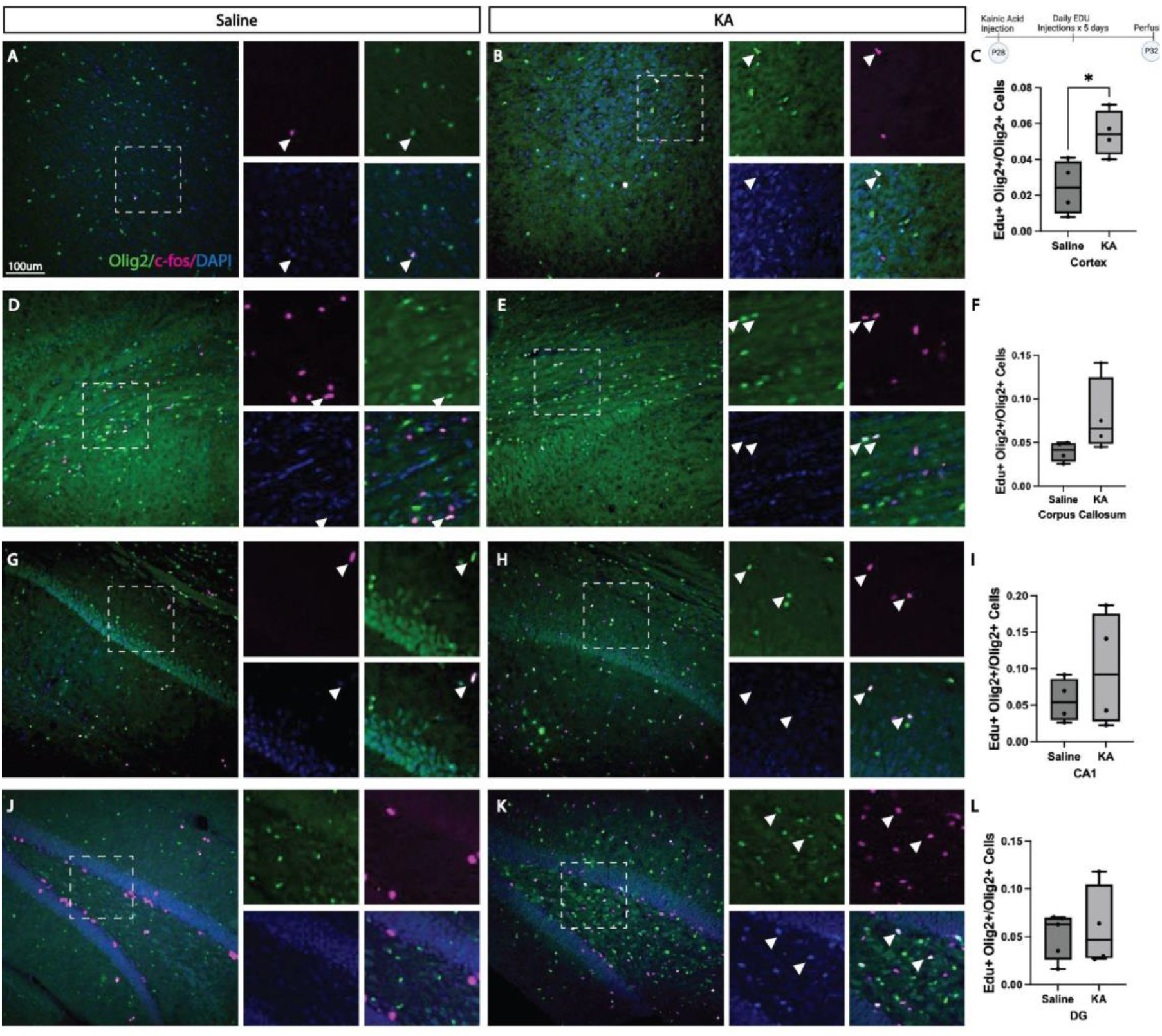
Kainic Acid Induced Seizure Leads to Increase in OPC Proliferation. Coronal sections from sex-matched littermates were stained using immunohistochemistry for Olig2 (green) and click-it chemistry for Edu (magenta). Control littermates received saline IP injections (A, D, G, and J), and experimental littermates received kainic acid IP injections up to 30 mg/kg (B, E, H, and K). Stains were visualized in the cortex (A and B), corpus callosum (D and E), CA1 (G and H), and DG (J and K). Boxes correspond to higher magnification images on the right; arrows indicate Edu+/Olig2+ cells, counterstained with DAPI (blue). Quantifications of Edu+/Olig2+ indicated in figures C, F, I and L for respective brain areas. C *unpaired student’s t-test, (t=3.073, df=6), p=0.0219. Scale bar, 100μm.

We then examined the overall pools of OPCs compared to mature oligodendrocytes, to determine if seizure activity affected both immature and mature oligodendrocytes equally and if all brain regions were affected. Interestingly, we found this to be variable by brain region, and by grey vs. white matter. We found that in the cerebral cortex, there was an overall decrease in the total density of Olig2+ cells in kainic acid-injected mice compared to saline injected mice, indicating a loss of oligodendrocyte-lineage cells as a result of adolescent seizure. Because Olig2 labels the entire oligodendroglial lineage, this loss could be due to a loss of OPCs, immature oligodendrocytes, or mature oligodendrocytes. We co-stained brains with Olig2 and CC1, and quantified the percentage of OPCs (Olig2+ CC1-cells) as well as the percentage of mature oligodendrocytes (Olig2+CC1+ cells) (Figure 3). We found no significant decrease in the density of mature oligodendrocytes in the cerebral cortex, but did find an increase in the ratio of OPCs to mature oligodendrocytes, indicating that oligodendrocyte-lineage numbers decrease and OPC numbers increase after seizure in the adolescent somatosensory cortex. In the corpus callosum, we found a significant decrease in the density of oligodendrocyte-lineage cells, but no significant changes in the density of mature oligodendrocytes or the ratio of OPCs to mature oligodendrocytes in KA-induced seizure mice compared to saline controls. In the CA1 of the hippocampus, we found that the density of oligodendrocyte-lineage cells increased, even as the density of oligodendrocytes decreased after seizure induction with KA compared to saline controls. The ratio of OPCs to mature oligodendrocytes increased in CA1 in mice after seizure, compared to saline controls. In contrast, in the dentate gyrus, we found no significant differences in oligodendrocyte lineage density, oligodendrocyte density, or the ratio of OPCs to oligodendrocytes in mice after seizure compared to saline controls.

**Figure 3.**
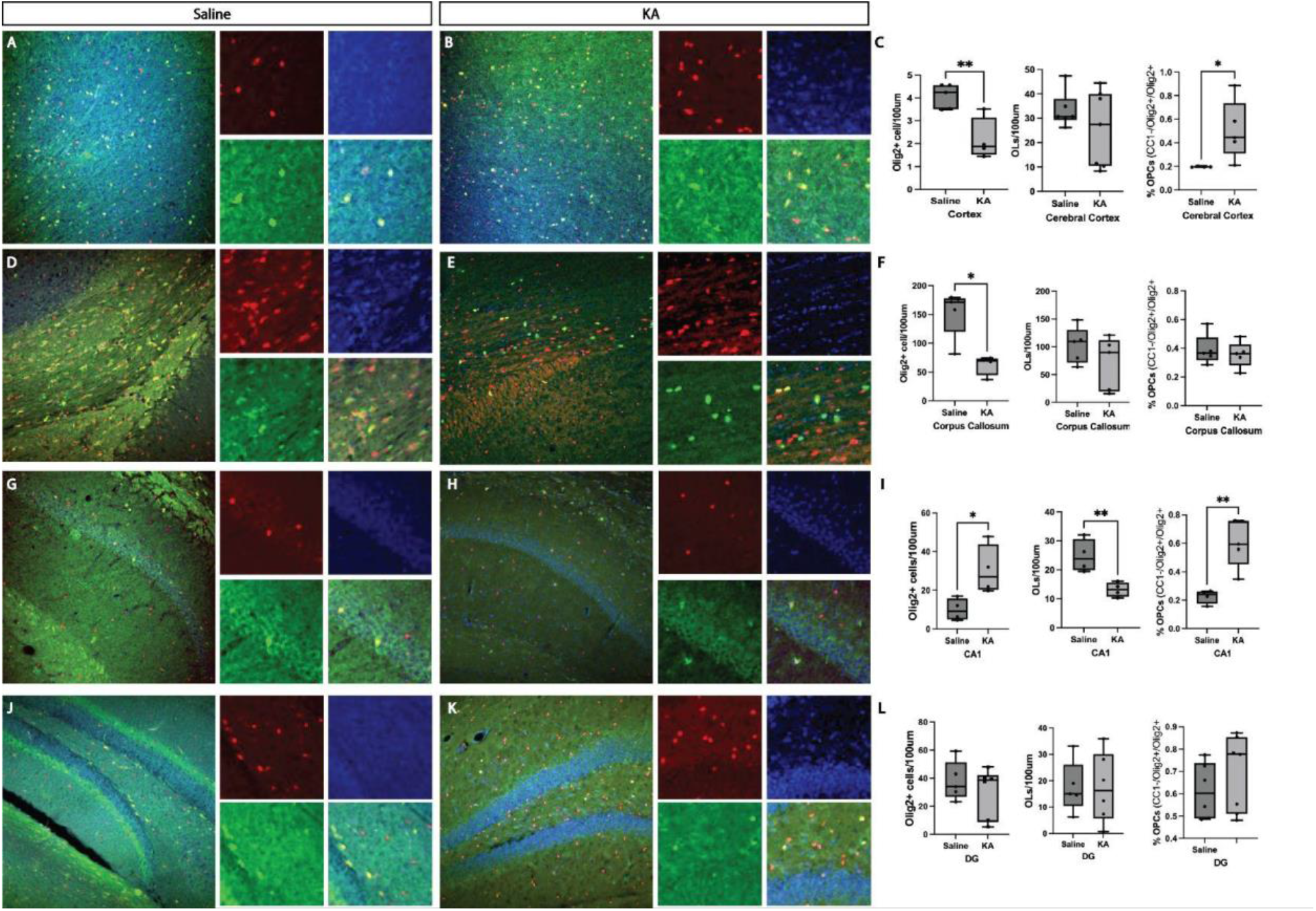
Kainic Acid Induced Seizure Leads to an Increase in OPCs. Coronal sections from sex-matched littermates were stained using immunohistochemistry for Olig2 (green) and CC1 (red), and counterstained with DAPI (blue). Control littermates received saline IP injections (A, D, G, and J), and experimental littermates received kainic acid IP injections up to 30 mg/kg (B, E, H, and K). Stains were visualized in the cortex (A and B), corpus callosum (D and E), CA1 (G and H), and DG (J and K). Boxes correspond to higher magnification images on the right. Scale Quantifications of Olig2+ cells, oligodendrocytes (OLs), and percentage of OPCs out of total oligodendrocyte-lineage cells indicated in figures C, F, I and L for respective brain areas. C * unpaired student’s t-test, (t=3.887, df=7), p=0.0060 for Olig2+ cells/100μm in Sal vs, KA, *unpaired student’s t-test (t=2.817, df=8) p=0.0226 for % OPCs/total Olig2+ cells in Sal vs. KA. F *Mann-Whitney test, (U=0), p=0.0159 for Olig2+ cells/100μm in Sal vs, KA. I *unpaired student’s t-test, (t=2.919, df=6), p=0.0267 for Olig2+ cells/100μm in Sal vs KA; *unpaired student’s t-test, (t=3.746, df=6), p=0.0096 for OLs/100μm in Sal vs KA; *unpaired student’s t-test, (t=4.272, df=7), p=0.0037 for % OPCs/total Olig2+ cells in Sal vs. KA. Scale bar, 100um.

Given that we saw changes in OPC proliferation, as well as the ratio of OPCs to mature oligodendrocytes, we wanted to investigate whether adolescent seizure impacted myelination. Seizure and epilepsy have previously been shown to cause demyelination, but this has not been tested after single seizure at adolescent development (Gibson, et al., 2018). We stained for MBP, and quantified the total myelin density by measuring the percentage of MBP fluorescence compared to total area. We found that in the cerebral cortex and in the corpus callosum there was a significant decrease in the area in which myelin was present in mice after seizure compared to mice after saline (Figure 4). We found that in the pyramidal layer, but not the stratum lacunosum moleculare of CA1, there was a significant decrease in myelin density in seizure induced animals compared to saline controls. Finally, we found no significant differences in the percent of myelin present between kainic acid-treated and saline treated mice in the DG of the hippocampus, either in the pyramidal layer or in the hillus.

**Figure 4.**
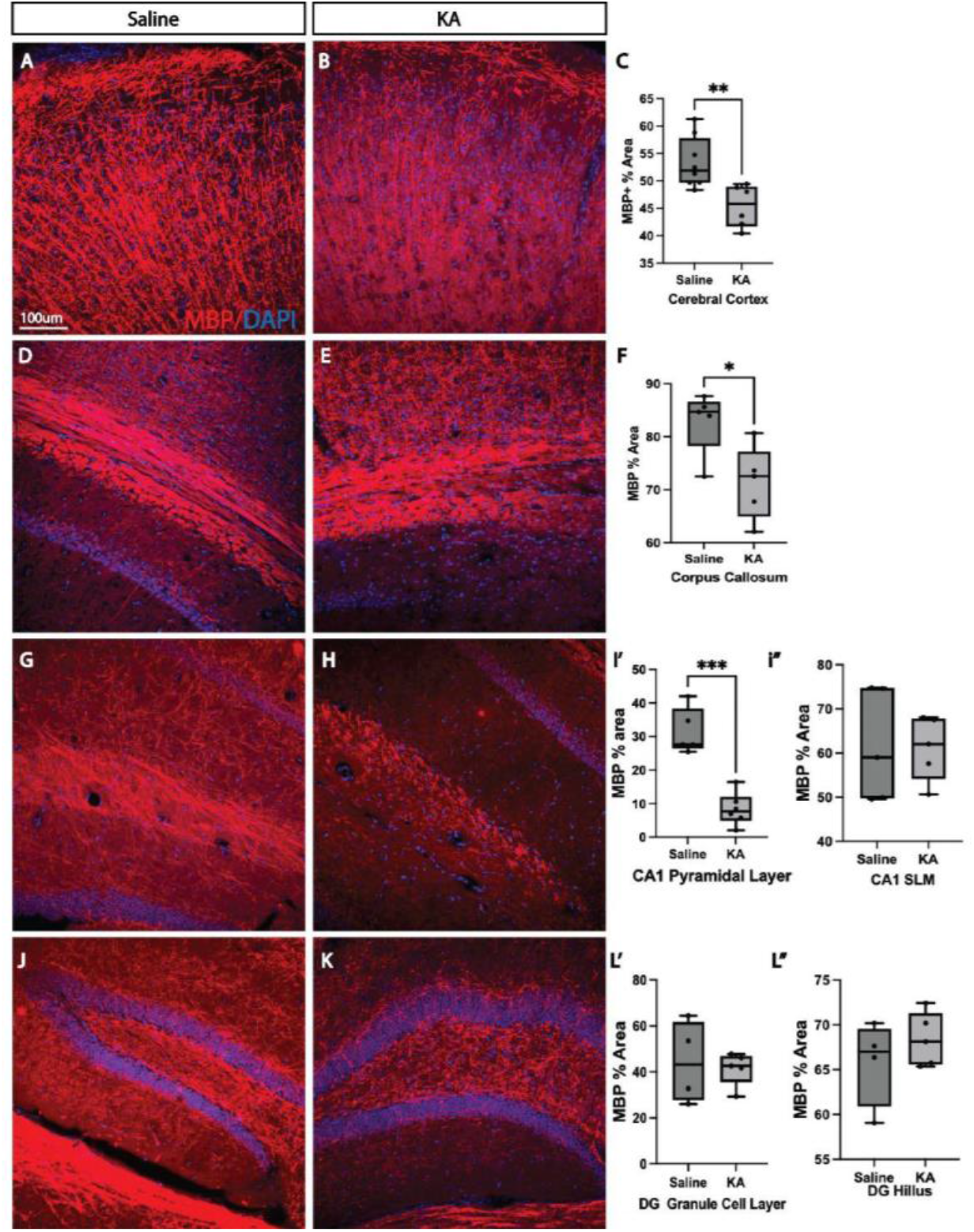
Myelination Decreases After Adolescent Seizure. Coronal sections from sex-matched littermates were stained using immunohistochemistry for MBP (red) and counterstained for DAPI (blue). Control littermates received saline IP injections (A, D, G, and J), and experimental littermates received kainic acid IP injections up to 30 mg/kg (B, E, H, and K). Stains were visualized in the cortex (A and B), corpus callosum (D and E), CA1 (G and H), and DG (J and K). Scale bar, 100μm. Quantifications of percent area covered by MBP+ cells indicated in C, F, I″, and L″ for respective brain areas, while percent area covered in the CA1 pyramidal layer and DG granule cell layer are indicated in I′ and L′ respectively. C *unpaired student’s t-test, (t=3.365, df=12), p=0.056) for MBP % area in Sal vs KA. F, *unpaired student’s t-test, (t=2.820, df=8), p=0.0225) for MBP % area in Sal vs KA. I’ *unpaired student’s t-test, (t=6.524, df=9), p=0.0001) for MBP % area in Sal vs KA. Scale bar, 100μm.

Considering the changes that we found in oligodendrocyte development and myelination, we wanted to examine whether seizure results in changes in the circuitry between neurons and OPCs. Previous work from our lab and others has shown the increased neuronal activity results in increased neuron to OPC connections, but given the pathological nature of seizure, which results in excitotoxicity, it was possible that these connections were lost as both neurons and OPCs suffered seizure related damage. We measured the number of neurons to OPC synapses using monosynaptic viral circuit tracing. We induced seizure using kainic acid injections, as explained previously, but on the fifth day, we injected pseudotyped deletion mutant rabies virus into the CA1 of the hippocampus in transgenic mice in which the proteins necessary for infection and spread were limited to OPCs. We waited 5 days for the virus to spread retrogradely from OPCs to connected neurons, then stained for neurons using Ctip2, and OPCs using Olig2. We quantified the number of starter OPCs and connected neurons using this approach, and found that the number of neurons connected to OPCs decreased in kainic acid-induced seizure mice, compared to saline injected controls (Figure 5). We confirmed that this was not due to variability in the number of starter OPCs (starter OPC numbers were not significantly different), indicating that the loss of neuron to OPC connections occurred as a result of seizure activity in adolescent mice.

**Figure 5.**
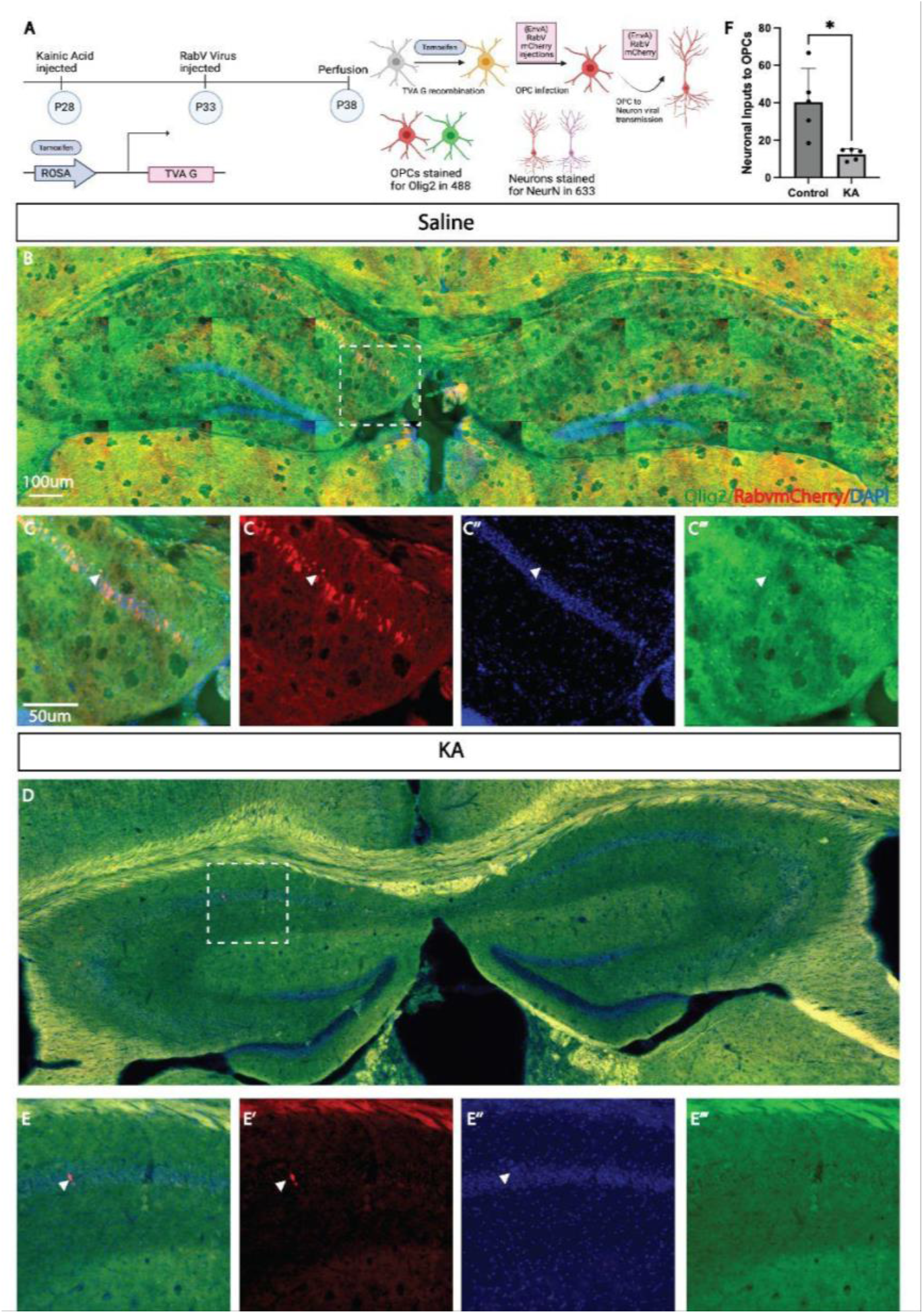
Neuronal Inputs Onto OPCs are Decreased After Adolescent Seizure. A. Schematic to illustrate transgenic strategy to target OPCs with TVA and G proteins necessary for infection and monosynaptic spread combined with(EnvA) SADΔGmcherry viral infection to retrogradely labeled neurons connected to OPCs. B. Connected presynaptic neurons are present in the pyramidal layer of CA1 in controls. C. Higher magnification optical sections illustrate initially infected (red+green) OPCs and connected neurons (red only). D. Connected presynaptic neurons are present in low density in CA1 and CA3 in the kainic acid-treated seizure mice. E. Higher magnification optical sections illustrate connected neurons (red only). F. Quantification of the number of presynaptic neuronal inputs onto OPCs in seizure mice compared saline controls. F *unpaired student’s t-test, (t=3.424, df=4), p=0.0245 for ratio of presynaptic neurons to connected OPCs in Sal vs. KA. Scale bar in B, 100um, Scale bar in C, 50um.

As seizure activity resulted in changes in myelin and neuron to OPC synapses in kainic acid-treated mice compared to saline controls, we wanted to determine whether seizure activity resulted in changes in ion channel expression in OPCs. We focused on the kir4.1 potassium channel, whose expression is linked to myelin function, and has been shown to be decreased after demyelinating injury (Song et al., 2018). As kir4.1 channel expression is visible on the somas of oligodendrocytes, we were able to determine whether there were changes in the global number of oligodendrocyte lineage cells that expressed kir4.1 by co-staining with Olig2 and Kir4.1. We found that Kir4.1 expression was significantly decreased in oligodendrocyte-lineage cells in the corpus callosum and CA1 of the hippocampus in kainic acid treated seizure mice compared to saline controls (Figure 6). We did not find significant differences in Kir4.1 channel expression in kainic acid treated seizure mice compared to saline controls in either the corpus callosum or dentate gyrus.

**Figure 6.**
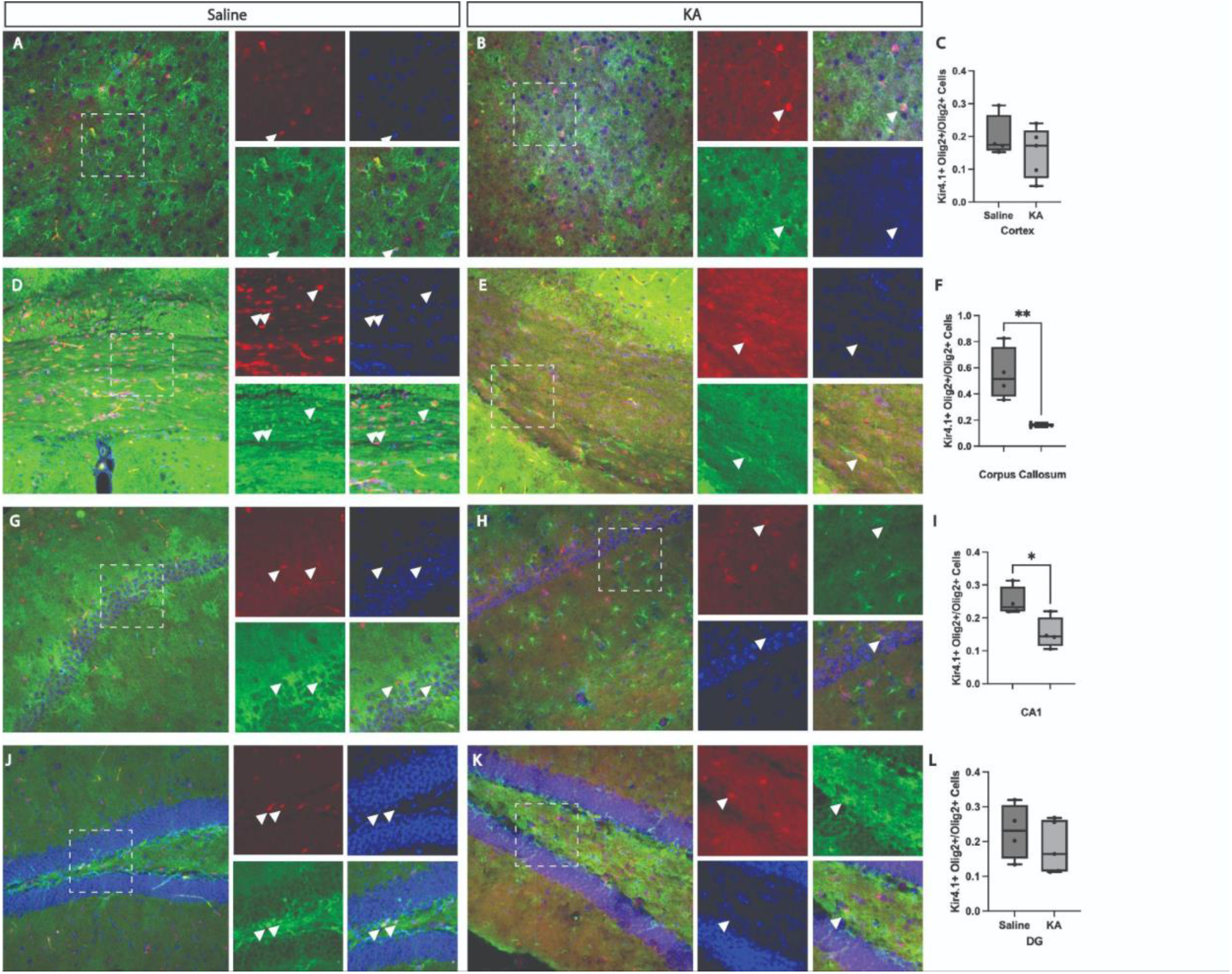
Potassium Channel Expression is Decreased in OPCs After Seizure. Coronal sections from sex-matched littermates were stained using immunohistochemistry for Olig2 (red) and Kir4.1 (green), and counterstained for DAPI (blue). Control littermates received saline IP injections (A, D, G, and J), and experimental littermates received kainic acid IP injections up to 30 mg/kg (B, E, H, and K). Boxes correspond to higher magnification images on the right. Scale bar, 100μm. Quantifications of Kir4.1+/Olig2+ indicated in figures C, F, I and L for respective brain areas. F *unpaired student’s t-test, (t=4.395, df=7), p=0.032) for Kir4.1+Olig2+/total Olig2+ cells in Sal vs KA. I *unpaired student’s t-test, (t=2.948, df=6), p=0.0257) for Kir4.1+Olig2+/total Olig2+ cells in Sal vs KA.

## Discussion

The time course of axon degeneration, cell loss, demyelination, and remyelination after injury is well understood, but the specific effects of seizure on OPCs in adolescents has not been previously tested. Given the frequency with which seizures occur in adolescents in general, and the number of children with various forms of epilepsy, understanding the changes in OPC development are important to understand the early changes occurring in the brain, with the goal of developing new therapeutic interventions. It is thought that dysregulation of either developmental or activity dependent myelination can contribute to status epilepticus onset and the progression to epilepsy, and that seizure itself can promote dysregulation of remyelination, increasing pathological circuit function that promotes further epileptic activity (Gibson, et al., 2017). Indeed, a recent paper has found that remyelination during epilepsy supported the recurrence of seizures, suggesting that remyelination is contributing to disease pathology over time (Knowles, et al., 2022). Our findings suggest that part of this mechanism may be due to the changes in neuron to OPC connections, which may be instructive for myelination, and are lost after seizure. Whether this loss of connectivity is due to the loss of neurons due to excitotoxic damage, or due to the loss of synaptic connections is not clear; this would be an interesting area of future explanation. The loss of kir4.1 potassium channels on OPCs is also an intriguing area for future exploration. Previous work has shown that loss of the kir4.1 channel in OPCs leads to spontaneous seizures, as this channel is important for regulating potassium levels in white matter (Larson, et al., 2018). This suggests a potential mechanism for the result that seizure induces aberrant myelination and promotes the development of epilepsy; gain of function experiments to rescue seizure as a potential preventative mechanism for the transition from status epilepticus to seizure would determine if Kir4.1 is important for this process.

Our results indicate that the OPCs of the dentate gyrus are largely unaffected by seizure. This is not surprising, as dentate gyrus neurons have changes in cell proliferation post seizure, but previous work has largely found defects in the CA1 region of the hippocampus. Due to the role of the entire hippocampus in seizure kindling, we chose to examine this region as well, although future studies would benefit from an analysis of the CA3 region as well, which was outside the scope of our study. In the future, we also propose that examining thalamic changes, as well as changes in the entorhinal cortex, would be important future directions to analyze additional networks known to be involved in seizure and epilepsy.

It is surprising that we found very different effects of seizure on the loss of oligodendrocyte lineage cells, as well as the number of OPCs and oligodendrocytes. In the cerebral cortex, we found that the total number of oligodendrocyte lineage cells decreased after seizure, but the number of OPCs increased and there was no significant loss of oligodendrocytes. Additionally, in the corpus callosum, the total number of oligodendrocyte lineage cells decreased after seizure, but the number of OPCs or mature oligodendrocytes was not different. Two explanations for these findings are possible. First, there may be high variability in the number of dying oligodendrocytes in the kainic acid treated animals. Our experimental design included animals with both stage 4 and stage 5 seizures, and we did not directly measure EEG activity during seizure induction. It is possible that there are larger differences in adolescent ages in the amount of oligodendrocyte cell loss that are responsible for this variability. A second possibility is that there is a loss in immature oligodendrocytes that is contributing to the total number of oligodendrocyte lineage cell decrease; this could be analyzed by examining the number of BCAS+ Olig2+ cells, which mark immature but not mature oligodendrocytes. We plan to follow up on this in future studies. Our results in the CA1 of the hippocampus suggest that while there is a net loss of cells from the oligodendrocyte-lineage cells, this likely can be explained by loss of mature oligodendrocytes, but may also be due to loss of immature oligodendrocytes (not measured here). We see a significant increase in the ratio of OPCs to oligodendrocytes, which is also consistent with this specific loss of mature oligodendrocytes, even as more OPCs are generated.

The changes in myelin area that we found present in the cerebral cortex, hippocampus, and corpus callosum are an area for further exploration. Given the plasticity of the adolescent brain, it would be interesting to examine if the remyelination that is likely to occur (at timepoints after our analyses) results in pathological circuit function and additional seizure activity, or is able to recapitulate developmental myelination and restore circuit function, preventing future seizures. This might also be mitigated by the timing of seizure induction at early developmental myelination timepoints vs. later time points where developmental myelination is being refined due to synaptic plasticity. Indeed, previous work has found that early establishment of temporal lobe epilepsy, at P12, if allowed to recover, results in restoration of myelin by three months of age, but it was untested whether the recovered myelin led to changes in circuit function consistent with abnormal remyelination (Bencurova, et al., 2022).

A number of developmental and neurological disorders have seizure comorbidity, including autism spectrum disorder, Down’s syndrome, multiple sclerosis, and auto-immune disorders such as lupus Lee, et al., 2015; Altuna at al., 2021; Kelley at el., 2009, Steriade, et al., 2021). Each of these disorders have abnormalities in glial cell function as well.

Understanding how the underlying developmental mechanisms of OPC development and myelination lead to symptoms in these diverse disorders and diseases may inform both our understanding of developmental and adaptive myelination in the healthy brain, as well as the changes that are due to OPC development, function, and connectivity after demyelinating injury or disease.

In conclusion, our work indicates that the impact of a single seizure has significant impacts on oligodendrocyte proliferation, survival, myelination, and circuit formation. Specifically, we found that seizure resulted in increased proliferation of OPCs in the cerebral cortex (Figure 2), increased proliferation of OPCs as well as a loss of mature oligodendrocytes in CA1, as well as more OPCs in corpus callosum, (Figure 3), and a decrease in total myelin in the cerebral cortex, CA1, and corpus callosum. Seizure also resulted in fewer neuron to OPC connections and a loss of Kir4.1 expression on OPCs, consistent with myelin damage. We plan to expand these analyses in the future to illuminate how synaptic proteins, including other ion channels, neurotransmitter receptors, and synaptic adhesion proteins are impacted during demyelination and remyelination in OPCs.

